# A digital DNA system reveals the superiority of unidirectional inheritance over ‘Lamarckian’ inheritance

**DOI:** 10.1101/2024.11.28.625825

**Authors:** Aswathi Shiju, Samantha D. M. Arras, Allen G. Rodrigo, Anthony M. Poole

## Abstract

In biology, changes to a DNA sequence can impact protein sequence but changes to protein sequence (phenotype) do not flow back into DNA (genotype). A system with bidirectional information flow (i.e. both translation and ‘reverse translation’) remains a theoretical possibility for an independent origin of life or an artificial biosystem, but the recent development of digital data storage in DNA does just this: changes made to a digital file can be written back into DNA, meaning changes to ‘phenotype’ can be written back to ‘genotype’. To explore the evolutionary properties of such a system, we created an artificial system where synthetic DNA serves as genotype and music as phenotype. Audio can be output from a DNA sequence, then recorded and written to DNA as ‘codons’, enabling bidirectional information flow (DNA→music and music→DNA). Our results show that the mutation rate in a bidirectional system is much higher than for unidirectional information flow, and that, under reverse translation there is no mechanism for preservation of codon choice across generations. This has the effect of eliminating the impact of spontaneous synonymous mutations, a key the benefit of a redundant genetic code. As a result, non-synonymous mutations are the only DNA-level changes that are transmitted across generations, and, as non-synonymous mutation can emerge at both ‘genotypic’ and ‘phenotypic’ levels, these occur at a two-fold higher frequency than in a unidirectional system. Our system holds some practical insight. First, for DNA read/write systems, it may be wise to avoid designing systems with ‘*de novo* reverse translation’ because the opportunities for mutation are higher; tracking genotype information from the preceding generation to guide this process may reduce error. Second, our system helps clarify how a ‘Lamarckian’ biological system might operate. We conclude that, were a ‘Lamarckian’ system of inheritance a feature of early genetic systems, it would likely have been short lived as the high frequency of mutation would risk driving the system to extinction. A system based on unidirectional information flow thus appears superior as there are fewer opportunities for mutational error.

## INTRODUCTION

All known biological systems exhibit a separation of genotype from phenotype. This separation likely evolved prior to the emergence of genetically encoded protein synthesis (1, 2). The advent of protein synthesis (3, 4) did however create a biophysical barrier between genotype and phenotype, a point first articulated in Crick’s Central Dogma of Molecular Biology (5, 6), where information is transmitted unidirectionally from DNA via RNA to protein, but not from protein sequence to either RNA or DNA sequence.

That said, there is ongoing interest in whether information transfer may flow from phenotype to genotype. Numerous biological systems have been speculated to show ‘Lamarckian’ features in that they appear to exhibit inheritance of acquired characteristics. Phenomena such as DNA methylation, RNA interference, horizontal gene transfer, and CRISPR-Cas-based adaptive immunity have all been suggested to be Lamarckian (7–12), though such claims have been questioned (13–17). Such speculation has a long history and is often attributed to Jean-Baptiste Lamarck (18), though this idea does not in fact originate with him (19).

Reverse translation, the transmission of information from protein to nucleic acid, has not been observed in natural systems. However, it has been argued to be biophysically possible (20–22) — a procedure for laboratory-based reverse translation has even been patented (23) — and reverse translation been suggested to have relevance to the early emergence of life (24). Another recently developed non-biological process also allows the concept of reverse translation to be probed: the use of DNA as a medium for digital data storage.

Digital information can be stored on synthetic DNA molecules (25–27), and is one of a number of competing alternative technologies (28, 29) to magnetic tape for digital archiving. In digital DNA storage, information can be stored in DNA, through conversion of data to a string of nucleotides. The exact sequence of nucleotides can be specified through synthesis using a DNA synthesizer, and DNA sequencing technologies enable that information to be recovered by reading the resulting DNA molecule (30). The conversion of DNA to digital information (e.g. a.txt file) is analogous to the process of gene expression, where a protein sequence is specified from the order of nucleotides in a DNA string. Information transmission in the opposite direction, from data file to DNA sequence, enables data to be written to DNA and is thus analogous to reverse translation. Thus, while phenotype to genotype transmission may not be a feature of modern biological systems, the use of DNA as a digital storage medium has this built into the read-write process, with digital data having even been directly written to the genomes of living cells (31–35).

We reasoned that a digital DNA storage system could be used to examine the properties of a system where phenotype to genotype information transfer is possible. As noted above, such a system is a theoretical possibility for an independent origin of life (either extinct, or existent elsewhere in the universe), or for an artificially developed biosystem. While it is tempting to ask the teleological question, ‘why has such a system not evolved?’, we limit ourselves to a tractable question: if such a system existed, how would it differ from known biological systems in maintenance and transmission of information? To probe this question, we created an artificial system that resembles the Central Dogma of molecular biology in that it possesses a genetic code, but where the information in the phenotype can be reverse translated to yield a new genotype. We reasoned that musical notation would be a good choice of phenotype as we could create a redundant genetic code (multiple codons per note) and we could easily assess phenotype for a musical score. Our code consists of 256 four-letter codons, coding for 64 musical note/duration combinations. The conversion table between music and codons can be used to convert from DNA to notes or vice versa. Using our system, we sought to assess the impact of changes to phenotype, the consequences of synonymous and nonsynonymous genotype mutations, and the impact of phenotypic changes on genotype. Our results reveal that spontaneous synonymous mutations are rendered irrelevant in a system with reverse translation, and that such a system has approximately twice the rate of evolutionary change when compared to a system that lacks reverse translation. Finally, we incorporated DNA synthesis, audio capture, and external selection pressure, to implement a functioning reverse translation system (**Figure 1**) that evolves in a manner that includes ‘Lamarckian’ as well as neo-Darwinian change.

**Figure 1.**
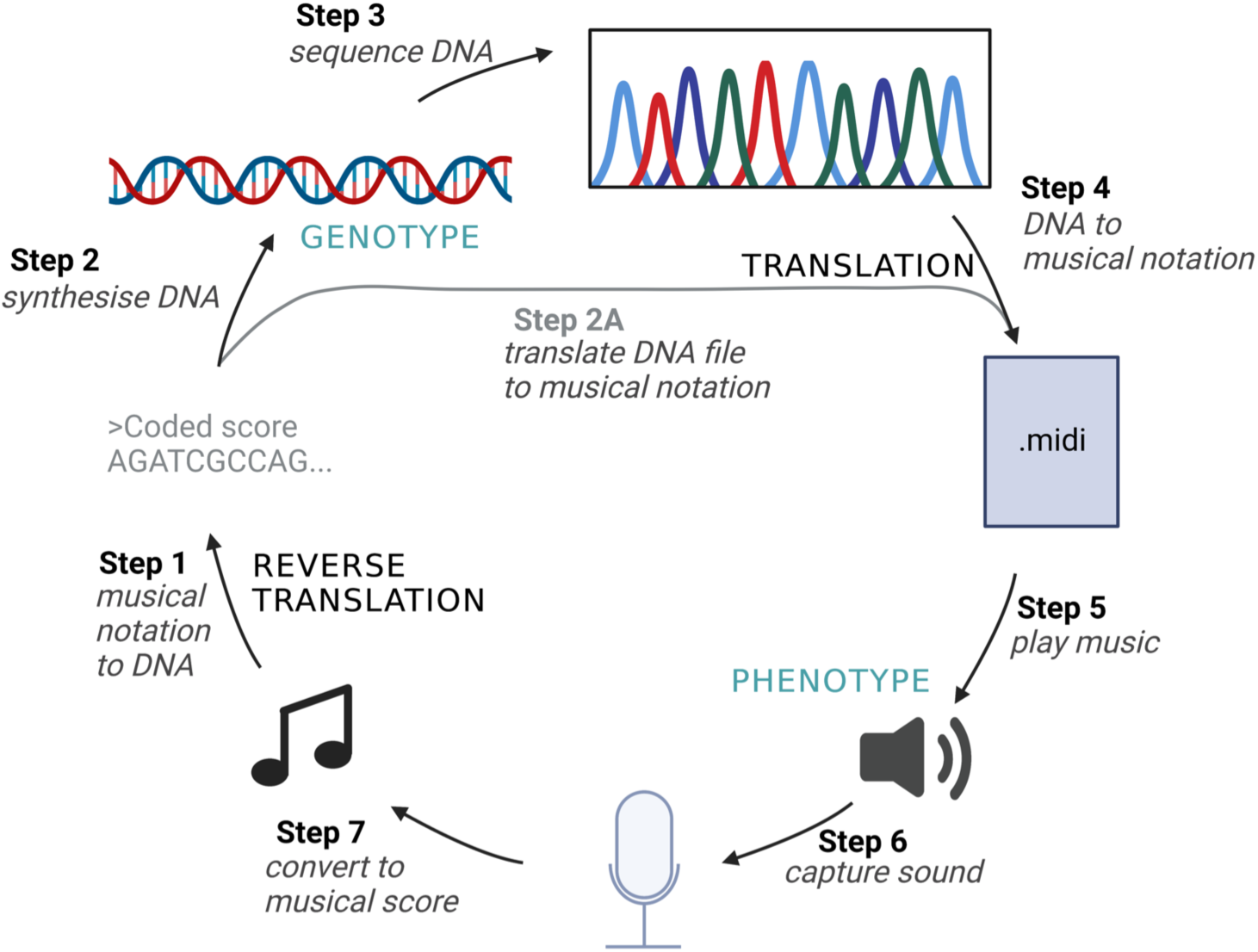
A digital system where phenotypic changes can be incorporated into genotype. In this system, sound is stored directly in DNA. A table that relates note pitch and duration combinations to 4-letter codons (see text for details) provides a means of converting between information stored as synthetic DNA (genotype) and sound (phenotype). It is possible to cycle between these two states, as follows. In Step 1, musical notation is converted to DNA sequence (reverse translation) using a conversion table that links musical notes and durations to DNA codons. The resulting DNA sequence file is used to guide synthesis of physical DNA (Step 2). The information within the synthetic DNA can be read by Sanger sequencing, generating a digital sequence file (Step 3). The resulting DNA sequence file is then ‘translated’ to musical notation using the conversion table, yielding a musical score in both MIDI and WAV formats (Step 4). Steps 2 & 3 can be skipped, allowing the DNA sequence file (FASTA) from Step 1 to be translated into musical notation (Step 2A, which is equivalent to Step 4). In Steps 5-7, audio is played through loudspeakers (Step 5), captured through recording (Step 6) and converted back into a musical score (midi file, Step 7). Created with Biorender.com

## METHODS

### Creation of a bidirectional inheritance system

In this section we describe the methods underpinning the seven steps depicted in **Figure 1**. Python source code created for this project is available at: https://github.com/PooleLab/Lamarck_DNA.

#### Step 1: Musical notation to DNA (reverse translation)

In this step, musical notes and durations are coded into DNA sequence. The artificial 4-letter genetic code consists of 256 codons, each coding for one of 64 musical elements. A musical element is a combination of a musical note with a specific pitch (depending on which octave it belongs to) and specified duration. We chose a subset of 16 notes from the MIDI standard, which comprises 128 notes (0-127) (**Table S1**). We created four durations for each of the 16 notes (1, ½, ¼, ⅛ beat), and used these to create a 4-letter code (256 codons) for 64 unique elements (note/duration combinations) (**Table S2**). A larger code would be required to accommodate all 128 notes plus a full range of durations, but a reduced set was sufficient for coding the music used in this project and served to constrain codon length and thus reduce DNA synthesis costs.

The input to step 1 is a musical score (**File S1**) composed of elements in MIDI format. This is converted to DNA sequence using the artificial code (**Table S2**). For each conversion of a musical element to DNA sequence, the corresponding codon is randomly assigned. The sequence is next checked for homopolymeric tracts of N=7 and, if a homopolymer is detected, the sequence is recoded. This check is performed to eliminate DNA sequences that could be challenging to physically sequence or chemically synthesize. The resulting sequence is output as a plain text file.

#### Steps 2 & 3: DNA synthesis and sequencing (read/write)

Text files from Step 1 (**File S2**) were sent for DNA synthesis (Step 2). The sequence of each synthesized DNA was confirmed (Step 3) by Sanger sequencing (**File S3**). DNA synthesis and Sanger sequencing was performed by Macrogen (South Korea). As these steps were performed by a commercial provider, data on background error was not available as the provider released only error-free data.

#### Step 4: Reading DNA and converting to musical notation (translation)

Our scripts accept either a plain text file with the DNA sequence as a string or a FASTA file. For experiments not involving DNA synthesis and sequencing (**Figure 1**, Steps 2 & 3; see results for details), the text file output from Step 1 was used as input for the conversion step. Where DNA synthesis and sequencing steps were used, a FASTA file was derived from sequencing data following completion of these steps; as data were error free (see preceding section), the raw DNA sequence derived from Step 1 and Step 3 were always identical.

The DNA file (plain text or FASTA format) is next ‘translated’ to yield a musical score. The decoder maps the 4-letter codons back to corresponding musical notes with distinct pitch and lengths, using the music code table (**Table S2**). The resulting musical score is then converted to MIDI file format (.mid) by appending each musical element along with the parameters for volume in dBfs (decibels relative to full scale), and tempo in bpm (beats per minute). We use-20 dBfs as the default value for volume (0.0 dBfs corresponds to the maximum value) and 100 bpm as the default tempo. An audio file in WAV format (.wav) is next generated from the MIDI file by first converting the notes to corresponding frequencies. The frequency (*ƒ*) of a note can be computed from the MIDI note number (*d*) as follows:

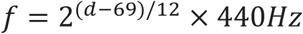

To capture note duration data for creation of the WAV file, the duration of each note, represented in ticks per beat in the MIDI file, is converted to ticks per millisecond, starting from the ‘note-on’ position to the ‘note-off’ position. Using the function Square() in pydub (https://github.com/jiaaro/pydub) (v0.25.1), tones are synthesized as a square wave.

#### Steps 5-7: Audio output, capture and conversion

The WAV file generated in the preceding step is played through loudspeakers as audio output (**Figure 1**, Step 5) either in a noisy environment (simulating high mutation during replication) or in a low noise environment (simulating low mutation), captured through a recording step (step 6), and converted into a MIDI file (step 7).

To capture audio and convert to MIDI format, we used Logic Pro (Apple, version 10.7.4, https://www.apple.com/logic-pro/). The WAV file from step 4 is played (step 5) and recorded (step 6) simultaneously, outputting a corresponding MIDI file (step 7). Depending on the signal to noise ratio during the audio capture, the resulting MIDI file may or may not be altered, compared to the MIDI file that the cycle began with (step 1, above). The output MIDI file can then be used as input for the next round of evolution.

### Pairwise comparisons for DNA sequences and musical scores

To trace mutational change in DNA over time in datasets where sequence length remained constant, Hamming distance was computed between a DNA sequence corresponding to each generation and the ancestor DNA sequence corresponding to the unmutated audio, using the following formula:

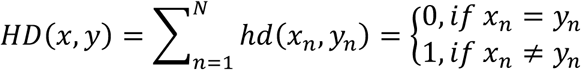

Where, for any two DNA sequences, *x* and *y*, with length *N*, Hamming distance *hd*(*x*_n_, *y*_n_) represents the number of different bases at the *n*^th^ position of the sequences *x* and *y*.

For comparison of DNA sequence datasets with varying length, we performed pairwise global alignments using Needle from the EMBOSS package (36) and default scoring. For pairwise global alignment of music scores, we developed a bespoke tool that made use of the Needleman-Wunsch algorithm, with the following scoring system: Exact-match (em) = 5, Near-match (nm) = 3, Mismatch (mm) = 0 and Gap penalty (gp) =-2. Exact matches are where both the note and duration are identical. A near match score is given where identical notes are of different durations. A mismatch score is given in cases where notes differ, regardless of duration. A gap penalty is applied for insertion or deletion states.

### Mutation and polling of versions of Ode to Joy

For the full cycle presented in **Figure 1**, five mutated versions of the melody “Ode to Joy” from Beethoven’s Symphony No. 9 were created. Version 1 was created by capturing sound (step 6) in a noisy environment. Version 2 was created by repeating steps 5-7 using Version 1 as input. Version 2 was used as input for creation of Version 3, and so on. This strategy was employed to create, at each round, a series of versions with different levels of mutation. Capturing sound in a noisy environment can lead to the generation of note/duration combinations that are not included in our code, which does not use the full range of notes available in MIDI. For elements that were not in our code, we chose the nearest equivalent. Our code does this by preserving the note but shifting the octave to the nearest octave specified by our code and either extending or shortening the duration to the nearest duration available.

The five versions from round 1 were then sent to our volunteer selection panel, each of whom independently chose a favourite/least disliked version. The version that received the most votes was converted to FASTA format and sent for synthesis and sequencing. This process was repeated for four rounds. In parallel, we performed a random selection process where 5 versions were created via the process described above, with one of the five versions chosen at random to seed the next round.

## RESULTS

To understand the properties of a system with bidirectional information transfer, we designed an inheritance system (**Figure 1**) where genotype is stored as DNA sequence and phenotype is music played over loudspeakers. This artificial system includes a DNA code where 256 4-letter codons correspond to 16 notes of four durations (**Table S2**). There is four-fold redundancy in this code, and it is grammarless (i.e., no start/stop codons). Our system differs from natural genetic systems because information transfer is bidirectional, i.e., information can be transmitted from genotype to phenotype (DNA→music) and from phenotype to genotype (music→DNA). As per the genetic code, changes at the level of genotype may impact phenotype, but only where mutations to the DNA sequence are nonsynonymous, i.e. leading to a change of musical element (**Table S2**). This process is labelled ‘translation’ in **Figure 1** (Step 4) and involves converting the DNA sequence into musical notation, with reference to the DNA code (a look-up table that enables conversion from 4-letter ‘codons’ to a corresponding musical element (i.e. a note/duration combination) (**Table S2**)). At the level of phenotype, alterations to the music can be transmitted across generations through a process we dub ‘reverse translation’ (**Figure 1**, Step 1). Here, musical elements are converted to DNA sequence, again with reference to the look-up table. Thus, any mutations to the musical score will lead to changes to the DNA sequence. Importantly, in reverse translation, the codons used in the preceding DNA sequence are lost once the musical notation has been created. Codons are instead randomly chosen from the look-up table during the reverse translation step. Finally, we include a short-cut (step 2A) which enables us to abrogate physical DNA synthesis and sequencing (steps 2 & 3) for the purpose of probing the behaviour of the system more rapidly using simulations.

To understand how mutations impact a system with bidirectional information transfer, we ran a series of six simulations each with a different mutational regime. We subjected a musical file containing Ode to Joy (the melody from the final movement of Beethoven’s Symphony No. 9; **File S1**) to one of six mutational regimes (**Table 1**) for 20 generations (where a generation is a full cycle of **Figure 1**, via Step 2A). For each mutational regime, ten replicate simulations were performed.

**Table 1.** Mutational regimes.

| Regime | Regime name | Actions at each generation |  |
| --- | --- | --- | --- |
|  |  | Step 1: Reverse Translation<br>(music→DNA) | Step 2A: Translation<br>(DNA→music) |
| 1 | 'zero' mutation | For each note/duration, select one **synonymous** codon<sup>a</sup> at random | Codons are converted to note/duration combinations with 100% fidelity |
| 2 | synonymous DNA-level mutation | For each note/duration, select one codon<sup>a</sup> at random, except for one codon, taken at random, where a **synonymous** codon is chosen<sup>b</sup> | Codons are converted to note/duration combinations with 100% fidelity |
| 3 | nonsynonymous DNA-level mutation | For each note/duration, select one codon<sup>a</sup> at random, except for one codon, taken at random, where a **nonsynonymous** codon is chosen<sup>c</sup> | Codons are converted to note/duration combinations with 100% fidelity |
| 4 | random DNA-level mutation | For each note/duration, select one synonymous codon<sup>a</sup> at random THEN make a single **point mutation** at a random position in the DNA sequence<sup>d</sup> | Codons are converted to note/duration combinations with 100% fidelity |
| 5 | nonsynonymous music-level mutation | For each note/duration, select one **synonymous** codon<sup>a</sup> at random | During translation, change one note/duration pair to a **nonsynonymous** <sup>c</sup> pair. All other codons converted to note/duration combinations<br>with 100% fidelity |
| **6** | nonsynonymous DNA-level<br>mutation plus nonsynonymous<br>music-level mutation | For each note/duration, select<br>one codon<sup>a</sup> at random, except<br>for one codon, taken at<br>random, where a<br>**nonsynonymous** codon is<br>chosen<sup>c</sup> | During translation, change<br>one note/duration pair to a<br>**nonsynonymous**<sup>c</sup> pair. All<br>other codons converted to<br>note/duration combinations<br>with 100% fidelity |

<sup>a</sup>Codons are drawn from Table S2
<sup>b</sup>All codons that specify the same note/duration pair differ by one nucleotide only. This action thus introduces a single synonymous point mutation.
<sup>c</sup>Nonsynonymous is defined as any note/duration pair that differs (by either note or duration) from the pair from the previous generation.
<sup>d</sup>Point mutation may result in either a synonymous or a nonsynonymous change.

The mutational regimes (**Table 1**) aimed to mimic natural mutational events. In conventional genes, the redundancy of the genetic code means that mutations to coding regions can be either synonymous or nonsynonymous. We therefore modelled the introduction of single mutations in DNA which were either synonymous, nonsynonymous, or random (meaning they could be either synonymous or nonsynonymous) (regimes 2-4). We included a ‘zero mutation’ control, where no DNA mutations are introduced (regime 1). This enables us to track the impact of the reverse translation step. For mutations to the musical score, there is no redundancy in that any change will alter the music score. However, as each note has four possible durations, it is possible for the note to be preserved but with a change to duration. Mutations can therefore impact pitch or duration, or both. For the purposes of understanding the mutational impact on the music score, we made no distinction between these variants, as they are all nonsynonymous, differing only in the degree to which the score is impacted (regime 5). We note that this is also the case for protein sequence differences; the fitness impact of nonsynonymous changes at the protein level can vary from neutral to lethal. Finally, we also modelled the impact of mutational change at both levels (DNA and musical score) because, in a system with bidirectional information transmission, mutations in both directions (translation and reverse translation; **Figure 1**) may occur (regime 6).

To track evolutionary change under regimes 1-6 (**Table 1**), we calculated the Hamming distance between the ancestral DNA sequence and descendants at each generation.

Hamming distances are suitable here because our regimes do not generate indels, and we simply wish to track the accumulation of mutations during each simulation. **Figure 2** shows the results of evolving our original Ode to Joy music file over 20 generations under the six regimes. After 20 generations, we observe no statistically significant difference in number of mutations between regimes 1 & 2 (P = 0.89, two-tailed t-test), despite regime 1 being ‘zero mutation’ and regime 2 including the introduction of a single synonymous mutation each generation (for a total of 20 mutations). This is because in our bidirectional system synonymous DNA mutations are not transmitted across generations—this information is lost in the act of translation, with reverse translation creating a DNA sequence by reference to the genetic code (21), not the DNA sequence of the preceding generation. This explains the large Hamming distance between ancestor and generation 1 sequences, because the latter are generated via an initial reverse translation event. As repeated reverse translation events mask synonymous mutations, the additional 20 mutations that are introduced at each round in regime 2 are eliminated and Hamming distances at generations 1 and 20 are not significantly different (P = 0.54, two-tailed t-test).

**Figure 2.**
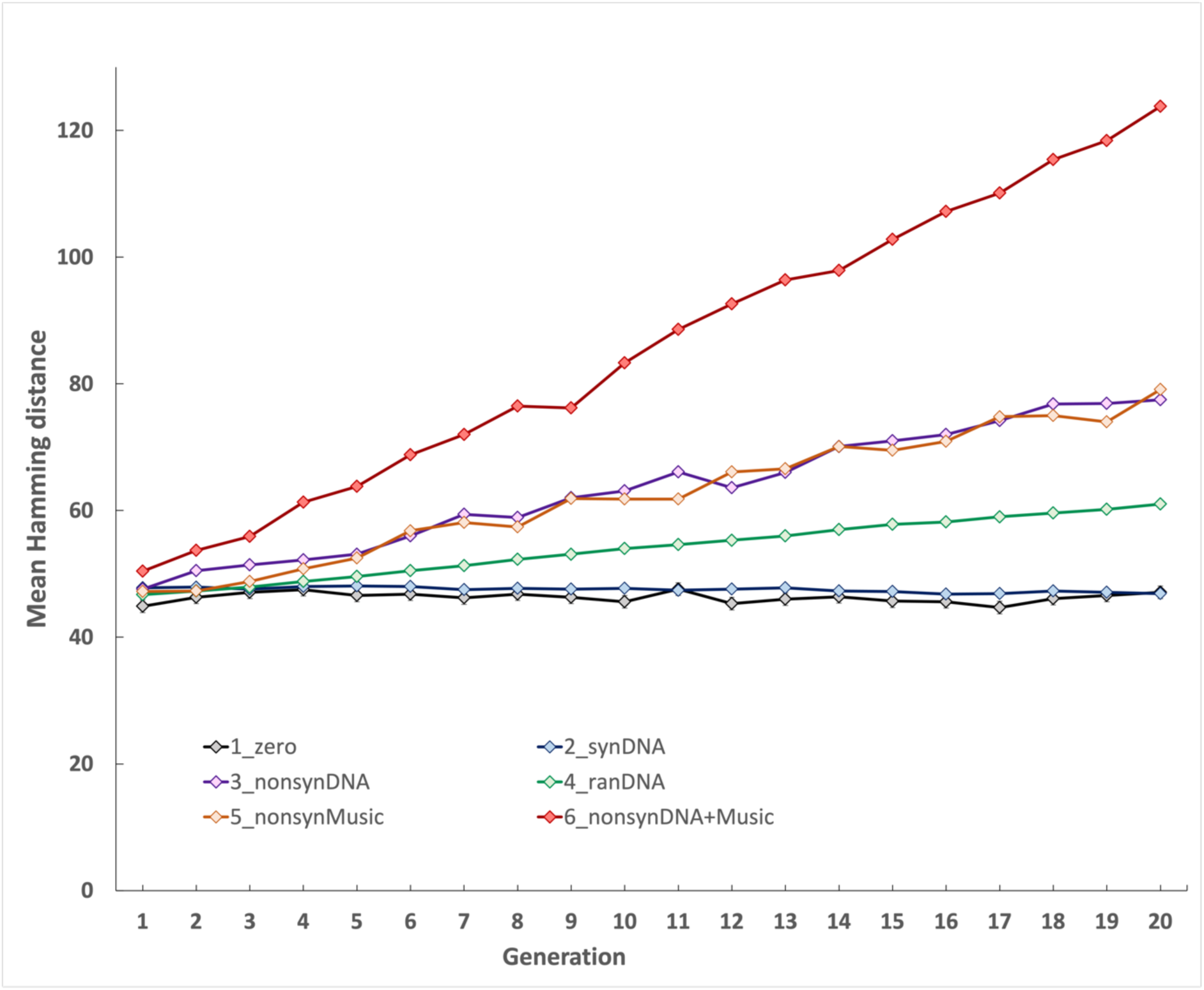
Comparison of mean Hamming distance between the ancestor and descendent generations under a range of mutational regimes. Each regime (Table 1) was iterated ten times across 20 generations, and Hamming distances were calculated at each generation for each replicate by comparison to the ancestor sequence. Regime 1: ‘zero’ mutation control (1_zero; black line); Regime 2: synonymous DNA-level mutation (2_synDNA; blue line); Regime 3: nonsynonymous DNA-level mutation (3_nonsynDNA; purple line); Regime 4: random DNA-level mutation (4_ranDNA; green line); Regime 5: nonsynonymous music-level mutation (5_nonsynMusic; orange line); Regime 6: nonsynonymous DNA-level mutation plus nonsynonymous music-level mutation (6_nonsynDNA+Music; red line). The x-axis shows generation number, y-axis shows mean Hamming distances for each regime calculated at each generation from comparisons between ancestor and the ten iterations.

By contrast, non-synonymous changes that occur either through mutation in DNA (regime 3) or in the musical score (regime 5) do result in an incremental increase in Hamming distance over time (**Figure 2**). By the end of the simulation, there is no significant difference in Hamming distances between these regimes (P = 0.34, two-tailed t-test) because the impact of mutation is equivalent, regardless of whether this occurs during translation or reverse translation. Regime 4 is intermediate between the synonymous and nonsynonymous regimes because random mutation of the DNA can lead to either a synonymous or a nonsynonymous change, with only the nonsynonymous cases resulting in an increase in Hamming distance relative to the ancestor.

Regime 6 results in much higher divergence from the ancestor (**Figure 2**) since in this ‘maximum mutation’ case, nonsynonymous changes occur at both DNA and musical score levels at each generation. While this example is extreme in in terms of mutation, it illustrates the additional mutational load in a bidirectional system; the maximal mutational impact for unidirectional transmission is given by regime 3 (20 nonsynonymous mutation events) which is approximately half that of regime 6 where there are 40 nonsynonymous mutational events under bidirectional transmission. Indeed, by 20 generations, the average Hamming distances between regimes 3 and 6 are significantly different (P = 1.15E-15, two-tailed t-test). Regime 4 is the most realistic for the DNA-level case (some mutations are synonymous, some are nonsynonymous) but this would only slightly reduce the mutational impact of bidirectional inheritance since all mutations during reverse translation would still be nonsynonymous. In summary, these results show that a system with bidirectional information transfer would be subjected to high mutation rates.

The above system allows a description of the impact of bidirectional inheritance on evolution. However, changes are neutral: change is recorded, but there is no consequence (either good or bad) associated with that change. To create something akin to an ‘environment’ that selects, we ran the full inheritance system from **Figure 1**, including DNA synthesis and sequencing, through four full cycles with environmental selection. We had hoped to include any errors at the DNA synthesis and sequencing steps, but using a commercial provider meant that we only received error-free data from these steps. To mimic environmental effects, we generated variants at Step 6, as follows. We performed sound capture (Step 6) in a high noise environment which resulted in the introduction of mutations (nonsynonymous substitutions, insertions, and deletions) to the musical score. For the resulting mutated score, we repeated steps 5 & 6 four further times to generate a series of five variants, each with successively more mutations. We then asked a panel of 13 volunteers to listen to the five audio files and select one file as their favourite/least disliked version. The version which received the most votes was converted to a musical score (Step 7) for the next round and the others were eliminated. Our panel thus served as the environmental selection (appeal to the human ear) over four rounds. To ensure they were not choosing based on our study aims, they were not provided information about the study design. One panellist with a background in music removed themselves from the polling for aesthetic reasons, stating that, “midi recordings have no musical qualities. All they are is a digital representation of pitch and duration with vapid timbre-there is no music in them”. Anonymised polling results for the remaining 12 volunteers at each round are provided in **File S4**. Music files for each round are provided for the interested reader to aesthetically assess for themselves (**File S5**).

To examine the impact of our selection regime, we also performed a parallel ‘neutral’ selection experiment where one of the five generated files was selected at random as the progenitor for the next round. We then compared the winners of each round to the ancestor. We performed two comparisons. First, we performed global pairwise sequence alignments between the selected DNA sequence from the current round and its progenitor (progenitor vs round winner). We then calculated the change in bitscore (Δbitscore) across rounds by subtracting bitscore from the bitscore derived from a comparison of the progenitor DNA sequence against itself (**Figure 3A; Tables S3 & S4**). Second, we created a pairwise global alignment tool (see Methods) to directly align musical scores, and tracked changes across rounds relative to the ancestor (**Figure 3B**). **Figure 3** shows that over the four rounds, our informal panel consistently picked the least mutated version of Ode to Joy (**Figure S1**), and thus the environmental effect resulted in purifying selection.

**Figure 3.**
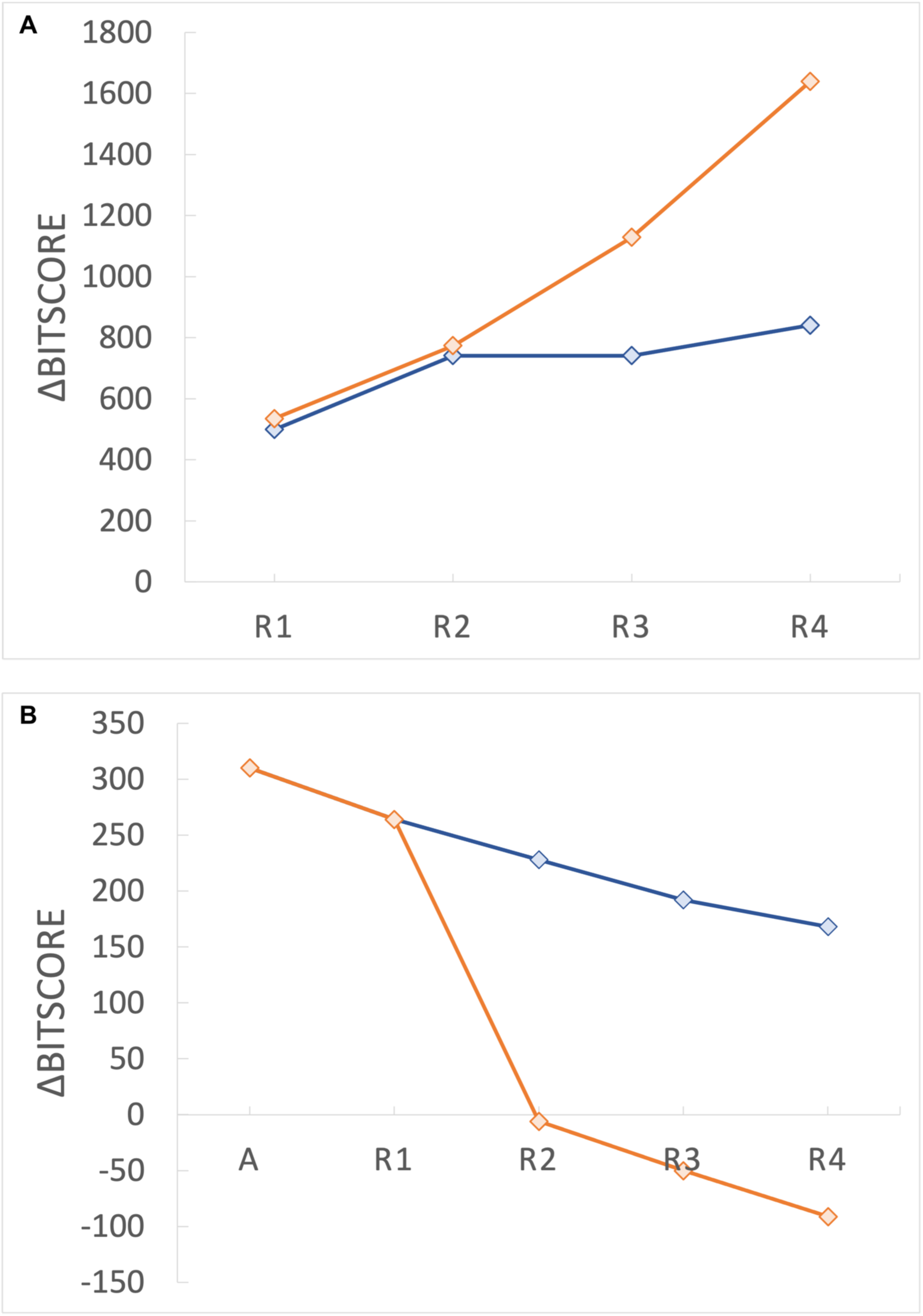
Effect of selection regime on evolution of DNA sequence and musical score. **A.** Bitscore comparison of DNA sequences selected by majority wins human poll (blue) versus random choice to mimic neutral evolution (orange) for four rounds of evolution (R1-R4). **B.** Bitscore comparisons between ancestor musical score (A) and human poll (blue) or random choice (orange) for R1-R4. The first data point represents the music-score comparison of ancestor against itself to establish the maximum similarity score (310).

## DISCUSSION

It has been noted that, with the benefit of a molecular level understanding of heredity, ‘Lamarckian’ inheritance of acquired characteristics would require information to flow in the opposite direction from that described by the central dogma of molecular biology, requiring a reverse translation step. In this conception, an environmental cue would generate changes to a protein sequence that would then need to be reverse translated from protein and reverse transcribed to DNA, resulting in an environmentally induced change to DNA. In such a system, changes in DNA would also impact protein sequence, so information transmission is bidirectional. While such a system has been theorised to be possible (20–22, 24), it is not a feature of natural biological systems. However, computing routinely makes use of read/write technology for the deposition, retrieval and editing of data. The development of systems for DNA-encoded digital data storage (25–27) thus makes it possible to create an artificial, non-biological system with some of the features expected for a biological ‘Lamarckian’ system. We report on the behaviour of just such an artificial informational system with bidirectional inheritance. Our system made use of 64 musical elements (16 musical note x 4 duration combinations), enabling us to mimic both reverse translation and the environmental inheritance of characters, with the latter provided through polling of volunteers on a series of musical files over four rounds of evolution.

*In silico* evolution of our system across 20 generations reveals that synonymous mutations are masked (**Figure 2**, schemes 1 vs 2) because there is no mechanism for their transmission across generations. This follows because reverse translation with a redundant DNA code (multiple codons for a single musical element) has no mechanism for intergenerational codon memory. In our system, a random codon assignment is made. This is consistent with hypothetical reverse translation systems (20) where there is also no mechanism to link a protein sequence to specific codons from which that protein was originally translated. Thus, our first conclusion is that a reverse translation mechanism renders synonymous mutation events irrelevant.

Our results also show that reverse translation increases mutation relative to a system without this process. As noted in methods, outsourcing synthesis and sequencing steps to a commercial provider meant we were unable to measure mutation rates of these processes at key steps within our system. However, we were able to track the impact of mutational opportunity by introducing mutations at each informational transfer event. As shown in

Figure 2, non-synonymous mutations can emerge at both DNA and musical element levels (scheme 6). For DNA, random mutations can be synonymous or non-synonymous (scheme 4), with only the latter contributing to intergenerational change (cf. schemes 4 and 2). As, reverse translation results exclusively in non-synonymous mutations (schemes 1 & 5), it creates a much higher potential for mutation than random DNA-level mutation (scheme 4) and is equivalent to a situation where DNA mutation only yields non-synonymous changes (scheme 3).

To introduce selection into our system, we polled volunteers who were given five mutated versions of Ode to Joy and asked to select their favourite/least disliked version (see methods). We found that this resulted in significant deviation from neutral evolution (as measured by Δbitscore for DNA (Figure 3A) and a similarity score derived from pairwise comparison of musical scores (Figure 3B; see methods)). We found that, despite recording in a noisy environment that yielded high levels of mutation (**File S5**), our human polling resulted in a clear signal of purifying selection (Figure 3). This shows that a selection regime that applies environmental pressure can impact evolutionary trajectory.

While the system we devised is not a biological system, it nevertheless serves to reveal some interesting features of such a system, were one to exist. It is difficult to determine what exactly constitutes a Lamarckian system, not least because the descriptions of such a system by Lamarck (18) predate our modern understanding of genetics and population variance. In the modern molecular context, one key feature would surely be the existence of reverse translation (37). Thus, transposed onto modern knowledge, a system with bidirectional information transmission can be considered to possess at least one Lamarckian trait. In our system, the inheritance of acquired characters occurs through changes to musical elements during recording resulting from an environmental effect (a high noise environment), with changes being written back into DNA. While this is analogous to reverse translation, it creates two levels of genetics, one at the level of DNA and one at the level of music. DNA in our system is entirely genetic (there are no phenotypic characteristics), while the musical score is both genetic and phenotypic.

A second feature of Lamarckian evolution is its goal-directed nature (17, 37). In our system multiple lineages are generated, and we contrived to create selection between them, via human polling. While we applied a selection pressure through our human poll, the participants did not have an explicit, shared ‘goal’ in mind (they were not in communication), and given that our system involved selection between individuals with only one individual reproducing in the next generation, the system has a distinctly Darwinian flavour. The process of selection on standing variation is not an insight that was understood in Lamarck’s time. Hence, in our system, the differential survival of one version of five (the fittest variant), looks Darwinian. One might argue that the tendency of our participants to make selections that minimised mutation of a recognisable piece of music could imply goal-directedness, but our system still requires selection from a range of individuals. Thus, despite the presence of reverse translation, the system still exhibits the Darwinian traits of natural selection, and descent with modification. This can however be said to hold for all modern phenomena that have led researchers to suspect existence a Lamarckian evolutionary process. It is our understanding that none of those researchers are denying the existence of Darwinian evolution, they are simply noting that some aspects of the evolutionary process bear a resemblance to the process of evolution of acquired characters.

Our system explicitly requires this latter process through reverse translation (step 1), so fits a modern definition of a Lamarckian process.

With the emergence of digital data storage, and the capacity to read and write data in DNA in living organisms, it may be that a portion of the genomes of these new species could be considered to be capable of evolving in a ‘Lamarckian’ way. However, as noted above, this is far from mutually exclusive of Darwinian mechanisms of evolution, and a Darwinian view might be that insertion of digital data into DNA is simply another special case of recombination. It has been noted that horizontal gene transfer does not require novel Lamarckian mechanisms to be invoked in order to understand this process (14). One might call a system with reverse translation Lamarckian because changes to phenotype feed back into genotype. However, our system is also a two-level genetic system, blurring the genotype/phenotype distinction. Thus, while it is fair to label reverse translation as ‘Lamarckian’, it is still worth noting this process is quite far removed from what Lamarck understood of inheritance in 1809.

## AUTHOR CONTRIBUTIONS

Conceptualisation: AMP, AGR

Data curation: AS, AMP

Formal analysis: AS, AMP

Funding acquisition: AMP, AGR

Investigation: AS, SDMA, AMP

Methodology: AS, AMP

Project administration: AMP

Resources: AMP, AGR

Software: AS

Supervision: AMP, AGR

Validation: AS, SDMA, AMP

Visualisation: AS, AMP

Writing – original draft preparation: AS, AMP

Writing – review and editing: AS, AMP, AGR

All authors read and approved the final version of the manuscript.

## ACKNOWLEDGEMENTS

We gratefully acknowledge financial support of the Digital Life Institute and the School of Biological Sciences at the University of Auckland. The following individuals are gratefully acknowledged for their participation in the file selection process: Zacharias Campbell, Jason Chen, Michael Goldwater, David Kelley, Soon Lee, Sylvie Herman-Le Denmat, Emily Parke, Jasper Perry, Saskia van Rijnsoever, Nobuto Takeuchi.

## List of Supplementary materials

**Figure S1.** Music score comparison between the mutated audio files against the precursor audios using modified version of Needleman-Wunsch similarity calculation for the four-rounds.

**Table S1.** MIDI standard

**Table S2.** 4-letter DNA code for 16 notes x 4 durations **Table S3.** Bitscore calculations for purifying selection. **Table S4.** Bitscore calculations for neutral selection **File S1.** Ode to Joy score in MIDI notation (.txt).

**File S2.** DNA text files from Step 1 (.fasta)

**File S3.** Sanger sequencing chromatograms aligned to DNA files (.pdf)

**File S4.** Questionnaire and voting data (.docx)

**File S5.**.zip archive containing Ode to Joy ancestor and rounds 1-4 music (.wav)

